# Modifications to the SR-Rich Region of the SARS-CoV-2 Nucleocapsid Regulate Self-Association and Attenuate RNA Interactions

**DOI:** 10.1101/2023.05.26.542392

**Authors:** Patrick N. Reardon, Hannah Stuwe, Sahana Shah, Zhen Yu, Kaitlyn Hughes, Elisar J. Barbar

## Abstract

The nucleocapsid protein (N) of SARS-CoV-2 is essential for virus replication, genome packaging, and maturation. N is comprised of two folded domains that are separated by a highly conserved, disordered, Ser/Arg-rich linker, and flanked by disordered tails. Using NMR spectroscopy and analytical ultracentrifugation we identify an alpha-helical region in the linker that undergoes concentration dependent self-association. NMR and gel shift assays show that the linker binds viral RNA but this binding is dampened by both phosphorylation and a naturally occurring mutation, whereas in contrast, RNA binding to the full-length protein is not affected. Interestingly, phase separation with RNA is significantly reduced upon phosphorylation but enhanced with the mutation. We attribute these differences to changes in the linker helix self-association which dissociates upon phosphorylation but forms more stable higher order oligomers in the variant. These data provide a structural mechanism for how the linker region contributes to protein-protein interactions, RNA-protein interactions, liquid-liquid phase separation and N protein regulation.

## Introduction

Severe acute respiratory syndrome coronavirus 2 (SARS-CoV-2) is responsible for an ongoing global public health crisis, thus there is an urgent need to continue evolving our understanding of structures and functions of SARS-CoV-2 proteins and their role in the viral life cycle. The SARS-CoV-2 virion is composed of four structural proteins: spike (S), membrane (M), envelope (E), and nucleocapsid (N). N associates with the viral RNA genome, protecting and packaging it into the virion and it is also associated with the replicase transcriptase complexes (RTCs), which mediate the synthesis of genomic RNA (gRNA) in the virus-infected cell [1]. Because of these unique interactions, N plays critical roles in nucleocapsid assembly, virion assembly, mature virion packaging, and replicating and transcribing the single-stranded gRNA [2].

N is a 422-amino-acid long protein that is organized into two independently folded domains, the N-terminal domain (NTD) and the C-terminal domain (CTD). These domains are flanked by two disordered tails (N-IDR and C-IDR) and separated by a serine and arginine rich (SR-rich), largely disordered linker (Figure 1). The CTD forms a domain swapped dimer, facilitating dimerization of N [3] while the NTD is the primary RNA-binding domain [4], but both the NTD and CTD bind RNA [2, 5–9]. Our studies on SARS-CoV-2 have confirmed that the N protein is a dimer consisting of ordered and disordered domains and that the CTD is required for its stable dimerization [3]. The SR-rich linker is primarily disordered, with some evidence for a helix near the C-terminal end of the domain [10]. Subsequent studies have shown that the linker binds the ubiquitin like domain of nsp3a, adopting a helical conformation where it interacts with nsp3a, resulting in overall compaction of N [11].

**Figure 1.**
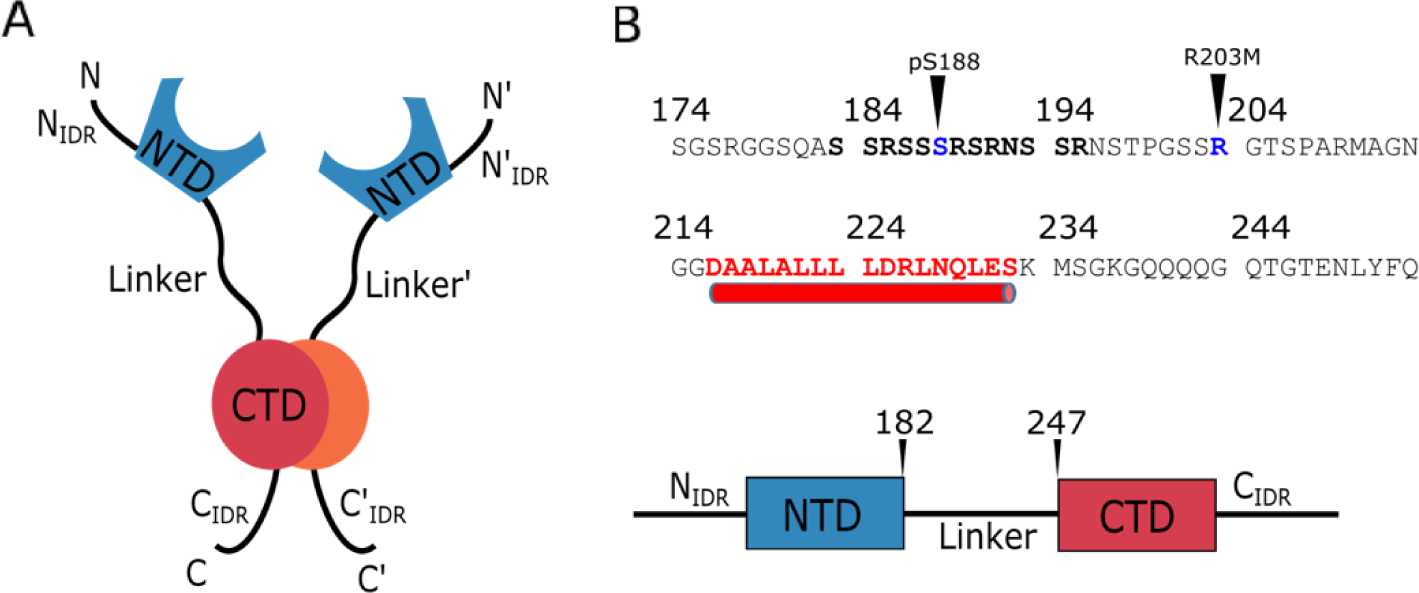
Overview of the nucleocapsid and the linker region structure. **A.** Schematic of the domain architecture of the nucleocapsid dimer. **B.** Sequence of the linker region with the SR-rich region shown in bold. The R203M and S188 phosphorylation site are indicated by arrows. The helical region is shown in red.

N carries out at least two distinct functions during the viral life cycle. First, it compacts and packages the genomic RNA in the virion. Second, it contributes to the formation of RTC’s. Both functions are associated with liquid-liquid phase separation (LLPS). Regulation of these functions is thought to depend on phosphorylation of N, with many of the phosphorylation sites occurring in the SR-rich region of the linker [12]. In addition, phosphorylation in the SR-rich region in SARS-CoV-1 has been suggested to alter host translation inhibition and modulate multimerization of N [13]. In SARS-CoV-2, phosphorylation of N altered LLPS, with unmodified protein forming gel-like condensates, while phosphorylated protein formed more liquid-like droplets [14]. The phosphorylation mediated differences in LLPS characteristics are hypothesized to contribute to switching between N functions [15]. Phosphorylation also alters the formation of particles in vitro that are similar to the ribonucleosome particles observed in the mature virion [16]. Overall, these observations demonstrate the critical role of phosphorylation in regulating the function of N during the virus lifecycle.

Over the course of the pandemic, several genetic variants of SARS-CoV-2 have emerged with increased infectious properties and are referred to as “variants of concern” [17–19]. Many of the mutations in N that are associated with variants of concern occur in the SR-rich region of the linker, suggesting that these mutations could alter regulation of N function and contribute to increases in virulence [20–22]. In a virus-like particle assay, some mutations in the SR-rich linker, particularly R203M, which occurs in the Delta mutant, dramatically increased the level of virus-like particle production [17], demonstrating that single mutations in N, especially those in the linker region, could alter and enhance viral function.

The above evidence suggests that the linker region of N is critically important for function and regulation. Yet, the detailed structure of linker and its various modifications, either through phosphorylation or mutation, have not been well characterized, nor has the functional contribution of linker and its modifications to RNA interaction or multimerization. Here we seek to fill in these knowledge gaps by directly probing the structure of WT and R203M linker proteins, as well as phosphorylated linker at serine 188, a known site of phosphorylation [12]. We also examine the interactions between the modified linkers and genomic RNA and compare to that of the WT. Our structural and functional characterization of the linker demonstrate how these modifications affect linker self-association, its interactions with RNA, and LLPS formation and provide a model of their role linker in regulation of N functions.

## Results

### Structural analysis of Linker N_175-245_

We used a suite of BEST triple resonance NMR experiments to assign the backbone resonances of Linker N_175-245_ at 10° C (Figure 2). Overall, the spectrum is consistent with a primarily disordered polypeptide, with generally poor chemical shift dispersion in the ^15^N-HSQC. However, we assigned a group of resonances between 7.6 ppm and 8.2 ppm, a region of the ^15^N-HSQC that is not typically associated with fully disordered polypeptide resonances, to residues 216-232 that exhibit carbon chemical shifts consistent with alpha-helical structure (Figure 2). Importantly, we assigned residues 225-230 that were missing from the previously reported linker assignments [10]. The chemical shifts for the remaining residues are consistent with random coil, indicating that the other regions of N_175-245_ do not adopt regular secondary structure under our experimental conditions. We assessed the fast time scale dynamics of N_175-245_ by measuring nuclear spin relaxation, ^15^N R_1_, ^15^N R_2_ and {^1^H}-^15^N NOE. {^1^H}-^15^N NOE for residues 178-215 and 236-248 were below 0.5 with negative NOE’s at the termini, indicating that the region is poorly ordered. ^15^N R_2_ relaxation rates in these regions were relatively uniform, with values between 2.5 s^-1^ and 5 s^-1^. ^15^N R_1_ showed relatively uniform relaxation with decreased values at the termini. As expected from our carbon chemical shifts, residues 216-235 exhibited elevated ^15^N R_2_ and {^1^H}-^15^N NOE, with NOE’s over 0.6, consistent with this region adopting secondary structure.

**Figure 2.**
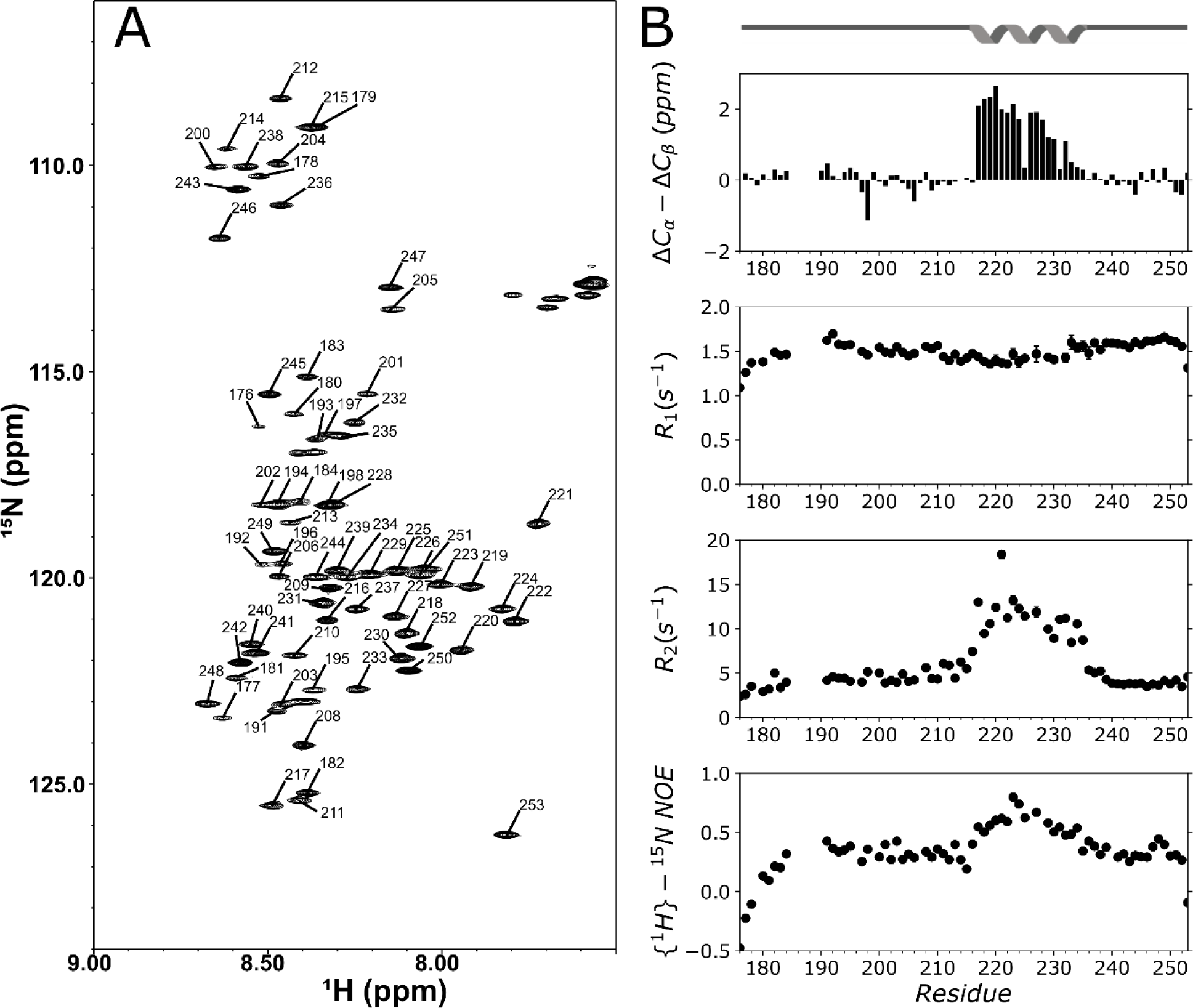
Structural characterization of WT N_175-245_. **A.** ^15^N-HSQC spectrum of 150 μM WT N_175-245_ at 10°C with resonance assignments. **B.** Chemical shift indexing and ^15^N nuclear spin relaxation with a cartoon depicting the location of the helical region (top of panel B).

While optimizing the conditions for NMR experiments, we noted that the spectrum at higher concentrations showed significant peak attenuation. At 300 μM, the N_175-245_ resonances corresponding to the helical region disappeared from the spectrum, suggesting that the helical region was self-associating (Figure S2). To confirm self-association in N_175-245_, we performed sedimentation velocity analytical ultracentrifugation (SV-AUC) with a GFP tagged N_175-245_ (WT-N_175-245_ GFP) to provide a stronger UV absorbance. The SV-AUC clearly showed two species in the WT-N_175-245_ GFP at 200 μM, corresponding to ∼3.1 S and 4.8 S, consistent with monomeric and dimeric (or possibly higher) species (Figure 3A). The GFP control at the same concentration only gave rise to a single peak, indicating that there is no detected dimerization of the GFP under these conditions (Figure S1).

**Figure 3.**
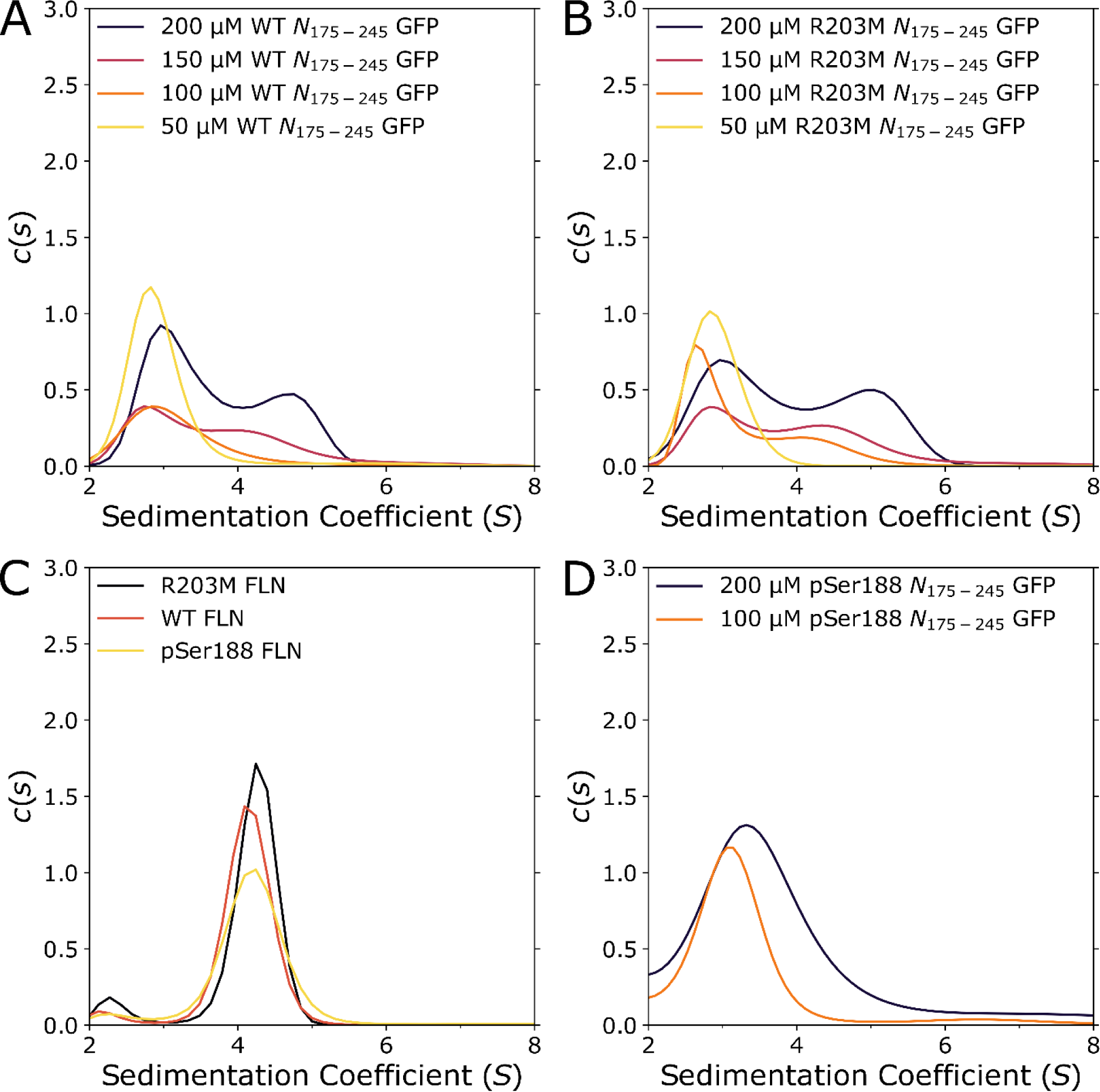
Sedimentation velocity analytical ultracentrifugation of N_175-245_ and FLN. All experiments with N_175-245_ proteins were performed with GFP fusion proteins on WT-N_175-245_ GFP at 50 to 200 μM concentration **(A)**, and R203M-N_175-245_ GFP at 50 to 300 μM concentration (**B)**. FLN WT, R203M and pSer 188 proteins at 22 μM concentration are shown in (**C),** and in **(D)** pSer188 N_175-245_ GFP at 100 and 200 μM concentration. GFP control at 200 μM concentration has a sedimentation coefficient of 3.1 (Figure S1).

### Structural analysis of R203M linker mutant

The R203M mutant of the SARS-CoV-2 Delta variant of concern enhances virion production in a virus like particle assay [17]. To determine if the R203M mutation alters the structure of the N protein linker, we analyzed R203M N_175-245_ using NMR spectroscopy and found that its ^15^N-HSQC spectrum was nearly identical to the WT protein, except for the site of the mutation. We also performed ^15^N R_1_, ^15^N R_2_ and {^1^H}-^15^N NOE analysis, which revealed little difference in the overall structure of the protein (Figure 4). Interestingly, what is different is that the R203M N_175-245_ mutant exhibited enhanced self-association when compared to the WT N_175-245_ protein. At concentrations above ∼45 μM the resonances corresponding to the helical region disappeared from the spectrum (Figure S2). In comparison, the helical region remained visible in the WT N_175-245_ protein at higher concentrations, up to 100 μM.

**Figure 4.**
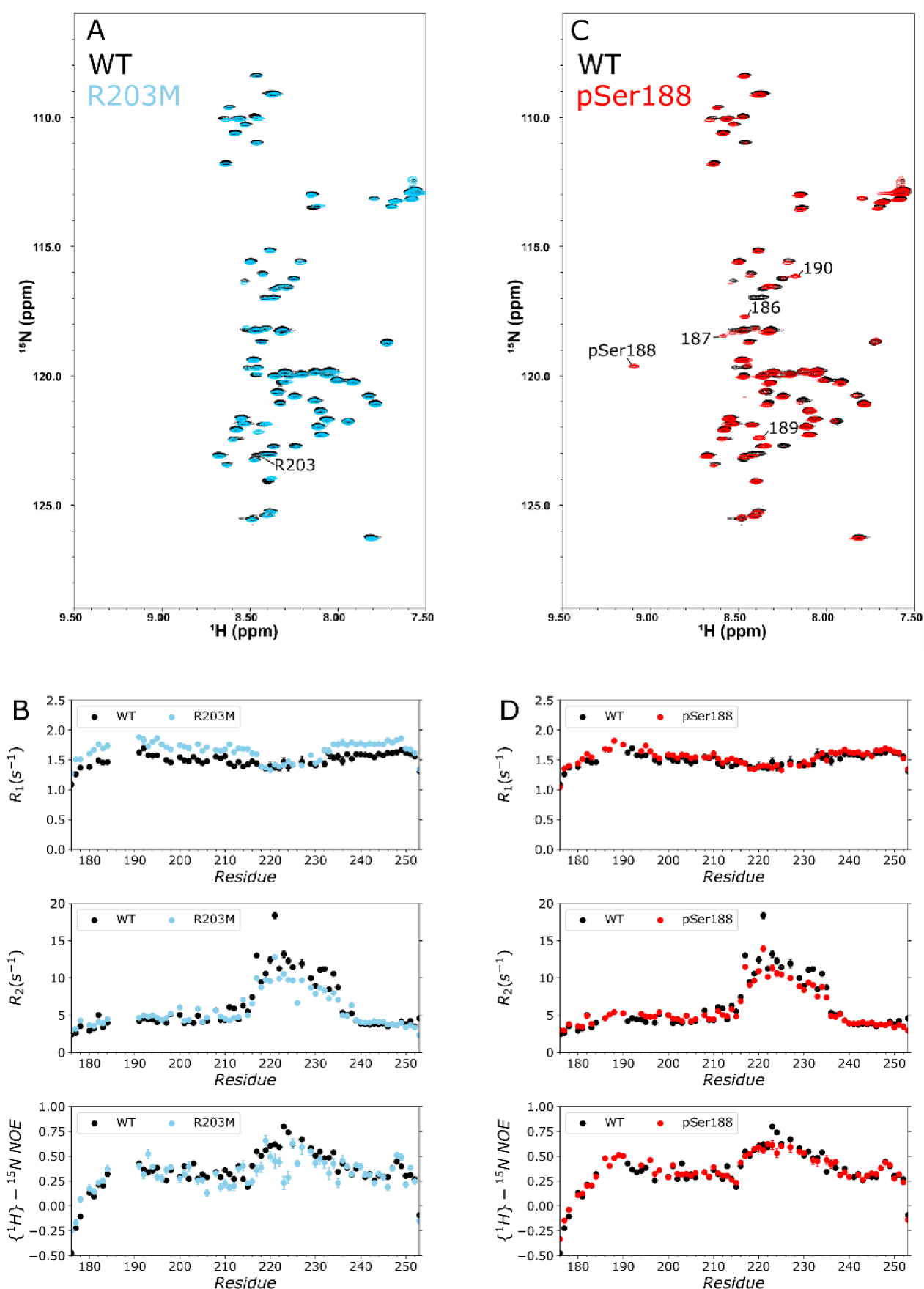
Structural characterization of R203M and pSer188 N_175-245_. **A.** Overlay of WT N_175-245_ (black) and R203M N_175-245_ (blue) ^15^N-HSQC spectra. **B.** Overlay of WT N_175-245_ (black) and R203M N_175-245_ (blue) ^15^N nuclear spin relaxation. **C.** Overlay of WT N_175-245_ (black) and pSer188 N_175-245_ (red) linker ^15^N-HSQC spectra. **D.** Overlay of WT N_175-245_ (black) and pSer188 N_175-245_ (red) linker nuclear spin relaxation.

To further determine the extent of self-association of R203M vs WT N_175-245_, we performed SV-AUC experiments (Figure 3). At 200 μM protein, R203M N_175-245_ gave rise to two species with sedimentation coefficients of ∼3 S and ∼5.1 S. These sedimentation coefficients were further separated from one another when compared to the WT N_175-245_ protein (∼3.1 S and 4.8 S), indicating slower exchange or possibly higher order complexes in the R203M. Estimating the area of each species in the c(s) plot indicates that 43% and 32% of R203M N_175-245_ and WT N_175-245_ are self-associated at a protein concentration of 200 μM. Lowering the concentration to 150 μM reduced the sedimentation coefficient for the second species in both R203M N_175-245_ GFP and WT N_175-245_ GFP, with the R203M N_175-245_ GFP second species running at a higher sedimentation coefficient compared to the WT N_175-245_ GFP. At 100 μM, the R203M N_175-245_ GFP still exhibited two species, while the WT N_175-245_ GFP showed only a single species, indicating that at 100 μM, WT N_175-245_ GFP is primarily monomeric while R203M N_175-245_ GFP still has a significant population of the larger species. At 50 μM, both WT N_175-245_ GFP and R203M N_175-245_ GFP showed only one species, consistent with primarily monomeric protein at this concentration. These results agree with our NMR observation that show resonance disappearance at lower concentrations for R203M N_175-245_ compared to WT N_175-245_, consistent with concentration dependent self-association. In all cases, the R203M linker showed increased sedimentation coefficients for the larger species vs the WT, indicating that the R203M N_175-245_ GFP has a stronger propensity to self-associate compared to the WT.

Having shown that the WT N_175-245_ self-associates in a concentration dependent manner and that the R203M mutation increases its self-association, we next determined if this observation holds true in the full-length proteins. With SV-AUC, R203M and WT exhibit similar sedimentation coefficients of ∼4.2 S, which is consistent with a dimer for FLN. We did not observe any higher order oligomers under these conditions. These results suggest that the self-association in the linker occurs within the same N dimer, rather than between N dimers to form larger oligomers.

### Structural analysis of pSer188 N_175-245_

Phosphorylation is thought to play an important role in regulating viral packaging and maturation [15, 23]. Since the residues of the SR-rich linker region are a major target for phosphorylation, we sought to determine if phosphorylation in this region alters the structure of the linker. We used genetic code expansion to incorporate a single phosphoserine at position 188 (pSer188), a known phosphorylation site. Comparison of ^15^N-HSQC of pSer188 N_175-245_ with WT N_175-245_ showed that the spectrum of pSer188 N_175-245_ was essentially identical to the WT N_175-245_, except for the resonances corresponding to residues proximal to the phosphorylation site. Comparison of ^15^N R_1_, ^15^N R_2_ and {^1^H}-^15^N NOE revealed that the proteins exhibited similar dynamic properties, with a modest increase in {^1^H}-^15^N NOE near the phosphorylation site (Figure 4). Phosphorylation typically results in a hydrogen bond between the phosphate and the backbone amide causing the strong downfield shift in the amide proton resonance. Formation of this hydrogen bond is expected to damp fast motion, so it is not surprising to see a modest impact on the {^1^H}-^15^N NOE in the region proximal to the site of phosphorylation.

We next determined if pSer188 N_175-245_ also exhibited self-association using SV-AUC analysis. Interestingly, SV-AUC analysis of pSer188-N_175-245_ GFP at 200 μM and 100 μM showed a single peak in the c(s) plots, with sedimentation coefficients of ∼3.3 and ∼3.1 S respectively (Figure 3). These sedimentation coefficients are similar to those observed for GFP alone (Figure S1). Importantly, the pSer188-N_175-245_ GFP proteins do not show a second distinct species at either 100 or 200 μM, indicating that self-association is considerably weaker for pSer188-N_175-245_ when compared to WT or R203M N_175-245_. The ^15^N-HSQC spectrum of pSer188 N_175-245_ at 300 μM protein also clearly showed peaks for the helical resonances (Figure S2), further supporting the conclusion that self-association of pSer188 N_175-245_ is considerably weaker compared to WT or R203M N_175-245_.

### WT, R203M, and pSer188 N_175-245_ interactions with RNA

RNA binding to the nucleocapsid is primarily driven by interactions with the N-terminal and, to a lesser extent, the C-terminal domains. However, more recent analysis has suggested that the linker could contribute to RNA binding [24]. Therefore, we determined if N_175-245_ can directly interact with RNA. We used an electrophoretic mobility shift assay to initially assess N_175-245_ binding to the first 1000nts of the 5’-end of SARS-Cov-2 viral RNA (g(1-1000)) [3]. The results show that N_175-245_ interacts with g(1-1000) RNA (Figure 5). Increasing concentrations of N_175-245_ caused an increasing shift of the g(1-1000) RNA to a higher molecular weight, indicating that the protein is binding to the RNA and reducing mobility in the gel in a concentration dependent manner.

**Figure 5.**
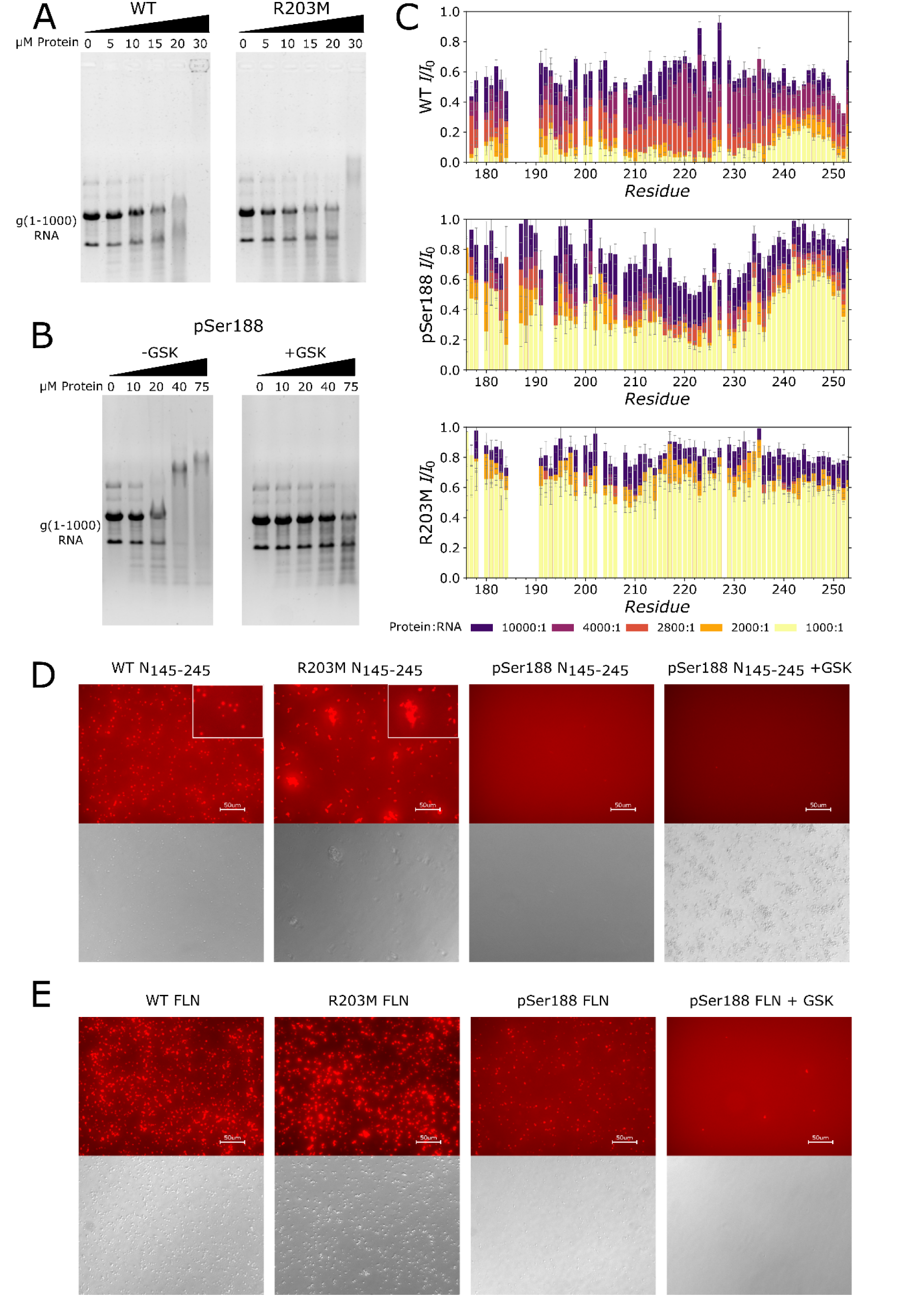
N_175-245_ and FLN interaction with g(1-1000) RNA and liquid-liquid phase separation. **A.** EMSA assay for WT N_175-245_ and R203M N_175-245_ protein (0-30 μM) and g(1-1000) RNA (0.5 μM). **B.** EMSA assay for pSer188 N_175-245_ (0-75 μM) with g(1-1000) RNA (0.5 μM).

We next used NMR to probe the RNA interaction site. Addition of increasing concentrations of g(1-1000) RNA to ^15^N labeled WT N_175-245_ caused most of the resonances to disappear, consistent with the formation of high molecular weight complexes. Essentially all of the resonances that are still observed following RNA addition, up to a 1000:1 protein:RNA molar ratio, are localized to the C-terminal region of the WT N_175-245_ and the residual TEV cleavage site (Figure 5). The rest of the protein, including the SR-rich region and the alpha-helical region, undergo significant line broadening indicating that the N-terminal part of the linker is the major RNA interacting site.

EMSA analysis showed that R203M N_175-245_ interacts more weakly with g(1-1000) RNA when compared to WT N_175-245_ where a modest increase in the concentration of RNA was required to shift the R203M N_175-245_ compared to the WT N_175-245_ (Figure 5). To further characterize this difference, we titrated g(1-1000) RNA into ^15^N labeled R203M N_175-245_ and observed that more RNA was required to significantly reduce the intensity of the R203M N_175-245_ resonances compared to the WT N_175-245_, indicating that binding of R203M N_175-245_ to RNA was weaker in comparison to the WT N_175-245_ (Figure 5).

GSK kinase was used to hyperphosphorylate (+GSK) the pSer188 N_175-245_ (-GSK). **C.** NMR titrations of g(1-1000) RNA into linker with a protein:RNA ratio varied from 10000:1 to 1000:1. Data were plotted as intensity ratios between with and without RNA resonances. **D.** N_175-245_ liquid liquid phase separation using fluorescence imaging (top) and bright field (bottom). Images were taken at 40X magnification and the scale bar is 50 μm. Insets are increased magnification to better show the differences in the particles. **E.** Similar to **D** but with FLN.

We next determined if phosphorylation altered RNA binding using pSer188 N_175-245_. We found that the pSer188 N_175-245_ also bound RNA, based on EMSA analysis, but the binding was somewhat weaker compared to WT N_175-245_ (Figure 5). To further characterize the interaction between g(1-1000) RNA and pSer188 N_175-245_, we titrated g(1-1000) RNA into ^15^N labeled pSer188 N_175-245_ and found that the pSer188 N_175-245_ exhibited RNA concentration dependent peak attenuation. As expected from the EMSA analysis, the binding was slightly weaker, with less peak attenuation at similar concentrations of RNA compared to the WT N_175-245_.

Having observed a modest impact on RNA binding for a single phosphorylation, we next examined the impact of hyperphosphorylation on RNA binding. The SR-rich region becomes hyperphosphorylated through the combined action of CDK1 and Glycogen synthase kinase 3 (GSK-3); GSK-3 phosphorylates serines or threonines four residues N-terminal of an initial site of phosphorylation. Phosphorylation at S188 would result in GSK-3 phosphorylating at positions 184, 180, and 176. Our EMSA results on GSK-3 kinase hyperphosphorylated pSer188 N_175-245_ show that hyperphosphorylation significantly reduced the interaction between N_175-245_ and g(1-1000) RNA because the linker showed no signs of interaction even at 75 μM, the highest protein concentration we tested.

### N_175-245_ RNA interaction in the context of FLN protein

Having demonstrated that the N_175-245_ linker alone has the capacity to interact with RNA and that this interaction is weakened by the R203M mutation, we next determined if these interactions are similarly affected in the FLN. We have previously shown that the disordered regions of FLN are observable with solution state NMR spectroscopy [3]. Therefore, we assigned several glycine resonances using our N_175-245_ assignments to probe RNA interaction by the linker in the context of FLN. Titration of g(1-1000) RNA into WT FLN to a final concentration of 1 µM resulted in attenuation of the resonances corresponding to glycines in the linker region, indicating that the linker also interacted with RNA in the context of the full-length protein (Figure S3). A similar titration with R203M-FLN showed that the linker glycine resonances were somewhat less attenuated with the same concentration of g(1-1000) RNA when compared to WT FLN. These results show that the linker interacts with RNA in the full-length protein and that this mutation alters this interaction, similar to what we observe with the isolated linker domains, even when the interaction with other domains remains intact.

### Liquid-Liquid Phase Separation is influenced by phosphorylation and mutation

Liquid-Liquid Phase Separation (LLPS) plays a role in viral maturation and is essential in packaging of the viral genome into the virus particle [15]. Given the reduced RNA binding affinity of R203M and pSer188 N_175-245_ relative to WT, we decided to examine the LLPS behavior with RNA with both N_175-245_ and FLN. WT N_175-245_ at 20 μM concentration phase separated at 37° C with 50 nM g(1-1000) RNA, showing spherical liquid like droplets. R203M N_175-245_ also formed particles under the same conditions as WT N_175-245_, however these particles exhibited less liquid like morphology. The particles formed by R203M N_175-245_ were fewer in number and larger on average than the WT N_175-245_ (Table 1). Interestingly, pSer188 N_175-245_ did not phase separate under these conditions, suggesting that phosphorylation inhibits phase separation.

**Table 1.** Count and average particle diameter for LLPS.

| Sample | Count | Average Diameter ($\mu\text{m}$) |
| --- | --- | --- |
| WT N<sub>145-245</sub> | 448 | 3.5 |
| R203M N<sub>145-245</sub> | 168 | 6.7 |
| pSer188 N<sub>145-245</sub> | 0 | N/A |
| pSer188 N<sub>145-245</sub> +GSK | 0 | N/A |
| WT FLN | 1080 | 3.3 |
| R203M FLN | 773 | 5.9 |
| pSer188 FLN | 402 | 3.2 |
| pSer188 FLN +GSK | 6 | 3.5 |

All experiments were performed with 50 nM g(1-1000) RNA. The N_175-245_ and FLN protein concentrations were 20 μM and 4 μM, respectively.

Having determined that mutation and phosphorylation can alter LLPS behavior of the N_175-245_ in isolation, the question remained whether similar results are observed in the context of the full-length N protein. Therefore, we examined the LLPS of R203M FLN, pSer188 FLN and WT FLN protein and compared their ability to form droplets to N_175-245_. Both WT and R203M FLN proteins phase separated with 50 nM g(1-1000) RNA at 37° C (Figure 5 and Table 1). The particles formed by R203M FLN protein were larger compared to WT FLN, but fewer in number. In contrast, phosphorylation reduced the propensity to phase separate under our conditions. The droplets formed with pSer188 FLN protein were on average similar in size to WT FLN, but fewer in number compared to WT and R203M FLN. The number and size of the droplets increased at 10 μM (Figure S4). The difference between WT FLN protein and phosphorylated N protein became even more pronounced when we hyperphosphorylated the pSer188 FLN protein using GSK-3. At 4 μM, the hyperphosphorylated pSer188 FLN showed almost no phase separation (Figure 5), with only a handful of droplets formed, clearly demonstrating that linker phosphorylation inhibits LLPS. Similar to the pSer188 FLN protein, the hyperphosphorylated pSer188 FLN showed increased LLPS at 10 μM protein (Figure S4). However, the number of droplets was less than the pSer188 FLN protein and much less than either R203M or WT FLN proteins.

## Discussion

### Self-association in the linker helix is modulated by phosphorylation and a neighboring mutation

The structure of N is generally depicted as having two folded domains that are linked together by a disordered linker and flanked by disordered regions at the N and C-termini. Our NMR chemical shifts and dynamics measurements confirm that the linker is primarily disordered but contains an α-helical region at residues 216-235. The helical region is C-terminal to the SR-rich region, which is thought to be important for regulation of N protein activity by phosphorylation and hyperphosphorylation.

Noteworthy about our experiments is that they were performed at conditions that allowed resonance assignments of the entire helix contrary to the deposited assignments which were missing in this region. Interestingly, we found that at protein concentrations over 300 μM many of the resonances corresponding to the helical segment would disappear, suggesting that self-association is the reason why these peaks were missing at the higher protein concentrations used in the published spectra. We confirmed self-association in the linker using SV-AUC, which clearly showed two species with sedimentation coefficients consistent with a mixture of monomer and dimer. Linker self-association could contribute to N protein dimerization, a function normally associated with the folded C-terminal domain, and also to higher order oligomerization, which might imply a role in genome packaging [19, 25]. In the context of the full-length N protein, we and others have also found that the resonances corresponding to the helical region are missing, suggesting a stronger self-association in full-length N [3, 11, 26]. Interestingly, the same region that forms a helix and self-associates in our study also forms a significant part of the interaction surface with nsp3a [11]. Based on our data, we propose that the linker-nsp3a interaction could be further regulated by the extent of self-association, with stronger self-association reducing nsp3a interaction.

SARS-CoV-2 has undergone rapid mutation over the course of the pandemic and a number of highly virulent variants of concern have emerged. Several mutations in the linker region of N are associated with these variants of concern, including Delta and Omicron. Mutations in the linker, most notably R203M, also enhance viral replication in a virus like particle assay [17]. Since the overall monomeric structure and fast time scale dynamics of the R203M mutant linker are essentially identical to the WT protein, the functional differences between the mutant and WT proteins are not attributed to their monomeric structures. Rather, since the R203M linker has a higher propensity to self-associate at the helical region, it is possible that part of the functional difference between WT and R203M linker is associated with the extent of dimerization of the linker helix. A recent report also shows that linker mutations can contribute to tetramerization of full-length N[19]. Based on our results, we suggest that the linker helical region is responsible for this additional self-association.

Both phosphorylation and mutations found in variants of concern fall within the SR-rich region of the linker, raising the possibility that mutation and phosphorylation could have related structural outcomes. Our results show that phosphorylation at serine 188, a site predicted to prime the linker for hyperphosphorylation by GSK-3, did not dramatically alter the monomeric structure of the linker. Similar to the R203M mutation, the functional differences between WT and pSer188 are not directly related to a change in the monomeric structure or fast time scale dynamics but to changes in self-association. Interestingly, whereas R203M increased self-association, the pSer188 linker exhibits reduced self-association compared to the WT and R203M linker proteins. We suggest that modulation of self-association in the linker region contributes to the mechanism for regulation of the N protein function by making the linker more available for interaction with other proteins, such as nsp3a, and reducing the formation of higher order oligomers formed by linker-linker interactions.

### Linker self-association modulates RNA binding and LLPS

While SR rich regions of proteins are often involved in RNA interactions [27, 28], direct experimental characterization of the interaction between SARS-CoV-2 linker and RNA has not been reported. Here we show that the linker interacts with viral RNA, in the absence of the other N protein domains, establishing that the individual NTD, CTD, and linker domain, are all capable of interacting with RNA with varying affinity and specificity. Further investigation will determine the relative contributions of each of these regions to N-protein RNA complex formation.

While the R203M Linker is structurally and dynamically similar to the WT, its reduced binding to RNA raises the question of how this reduction enhances viral packaging, when compared to the WT. Similarly, phosphorylation results in reduced RNA binding by the linker while hyperphosphorylation by GSK kinase, primed with pSer188, dramatically abolishes it. These results show that mutation and phosphorylation in the SR-rich region modulate RNA interaction and suggest some common mechanism for regulation of viral function. What makes this reduction in binding even more intriguing is that it is localized to the linker as full-length R203M, pSer188, and hyperphosphorylated pSer188 proteins still interact strongly with the RNA as shown by EMSA (Figure 5).

Phosphorylation of the linker by a combination of CDK-1 and GSK results in altered phase separation behavior, inducing a transition from gel-like to liquid-like droplets [15]. Our results confirm that hyperphosphorylation alters LLPS[15], but instead of conversion from gel-like to liquid-like droplets, we found that phosphorylation and hyperphosphorylation reduced the size and number of droplets formed, for both the linker and full-length proteins.

The R203M mutant also altered phase separation, with the morphology of the particles appearing more ‘gel’ or aggregate like compared to the WT protein. These results show that phase separation and RNA binding are distinct properties of N-protein. While RNA interaction is certainly required for phase separation, phase separation behavior can be modulated even if overall RNA binding remains similar. Modulation of phase separation appears to be at least in part regulated by the disordered linker region and its interactions with RNA, rather than only the folded domains. Finally, shifting to the self-associated linker could result in compaction of the RNA-nucleocapsid complex, preparing the complex for incorporation into the virus particle.

We propose a model for the linker contribution to N-protein structure and function (Figure 6). In the absence of RNA, the linker exists in equilibrium between open and self-associated states. Hyperphosphorylation and mutation in the linker region can shift this equilibrium in opposite directions. While linker RNA binding is reduced in the mutation and fully abolished in the hyperphosphorylated form, LLPS shows a different behavior. Hyperphosphorylation reduces linker interaction with RNA, reduces linker self-association and reduces phase separation, perhaps opening the linker for interaction with other proteins, while the rest of N remains bound to the RNA. On the other hand, mutations neighboring the linker helix promote self-association, and enhance the formation of aggregates in LLPS, which could contribute to compaction of the RNA-nucleocapsid complex. The linker must be freed from RNA or its self-associated dimer to interact with other binding partners during viral maturation such as the viral protein nsp3a [11] and the host protein 14-3-3[29]. Since the linker helix forms a major part of the interaction surface with nsp3a and is also the primary site of self-association, this suggests that modulation of self-association could also alter the interaction between nsp3a and linker. RNA interactions with linker could also alter interactions between nsp3a and N, by similarly blocking nsp3a from binding the linker. Interaction of linker with nsp3a induces considerable compaction of N that could be important for viral maturation [11], so weakening the interaction between linker and RNA may promote the interaction with nsp3a. Importantly, RNA binding by the NTD, and to a lesser extent the CTD, still drives strong interaction between nucleocapsid and RNA, so the nucleocapsid remains bound to RNA, but function is modulated by the linker modifications. Overall, this model suggests that regulation of N-protein interactions with RNA and other proteins is controlled by a combination of phosphorylation and mutations that act on the self-association of a critical linker helix in a disordered linker that is easily responsive to these site-specific changes and contribute to switching from genome replication to virus maturation.

**Figure 6.**
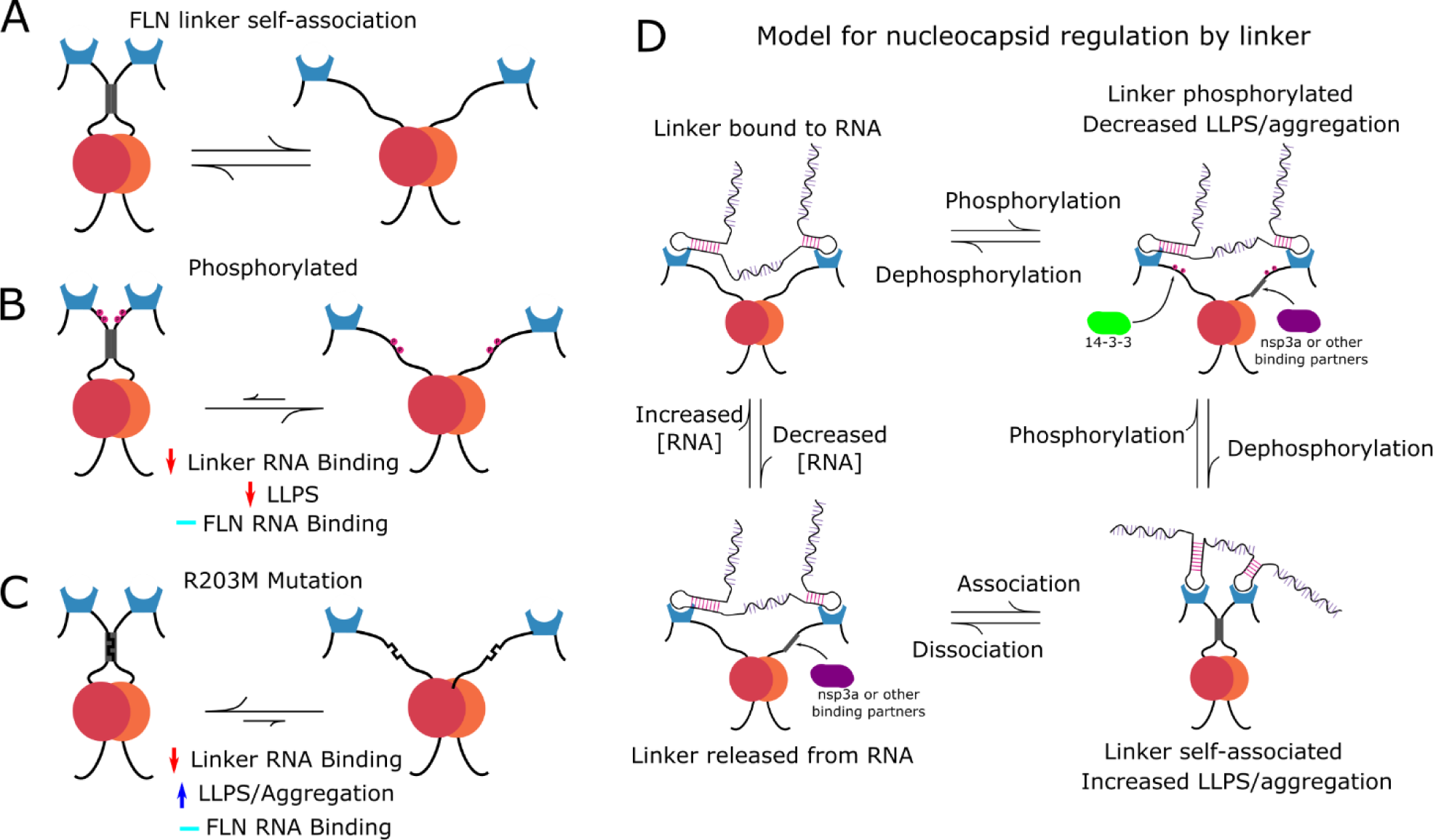
Model of phosphorylation and mutation effects on linker structure and function. **A.** Equilibrium for linker self-association at low concentration of nucleocapsid. Higher order oligomerization could be possible at higher concentrations. **B.** and **C**. Outcomes of phosphorylation and mutation. Phosphorylation reduces self-association, resulting in reduced linker-RNA interaction, reduced LLPS, but little change in FLN-RNA interaction because the NTD and CTD continue to bind RNA. R203M mutation also reduces linker-RNA interaction, but the increased linker self-association also increases LLPS and/or gel like aggregation. **D.** Schematic model showing the transitions between different RNA bound states of FLN. In the phosphorylated state, the linker region does not interact as strongly with RNA and it is not self-associated. It can interact with other proteins, such as 14-3-3 or nsp3a. At higher concentrations of RNA, linker can interact with RNA and is not self-associated. At lower concentrations of RNA, linker shifts towards the self-associated state and mutations can also influence this equilibrium and interaction with partner proteins.

## Materials and Methods

### Experimental design

Our goal is to examine the effect of phosphorylation and mutation in the SR-rich region in the linker connecting the two structured domains of SARS-CoV-2 N-protein. To this end, we prepared the linker and used NMR to identify an alpha helix secondary structure and prepared a phosphorylated and variant linker to identify their effect on the structure, and interactions with viral RNA. We also prepared the FLN protein and did similar experiments to show the effect of these modifications in the context of the full-length protein. We identified conditions at which the WT linker undergoes self-association on its own using analytical ultracentrifugation. The effect of these modifications on RNA binding were also followed by gel shift and droplet formation assays. Genetic code expansion methodology was used to insert a single phosphorylation site.

### Protein expression and purification

The N_175-245_ plasmid construct was prepared by inserting DNA encoding the linker region (amino acid residues 175 to 245) into a pRBC SUMO vector with a N-terminal bdSUMO tag, a C-terminal TEV protease cleavage site linked to GFP, and a hexahistidine tag [30]. The R203M N_175-245_ plasmid was prepared from the WT N_175-245_ plasmid with site directed mutagenesis (NEB site mutagenesis kit). The plasmids were transformed into BL21(DE3) *Escherichia coli*. To prepare unlabeled WT N_175-245_ or R203M N_175-245_ protein, BL21(DE3) *E. coli* with the appropriate plasmid were grown in Luria Broth (LB) rich media to an OD of 0.6-0.8. Protein expression was induced with 1 mM Isopropyl ß-D-1-thiogalactopyranoside (IPTG) at 37° C for 6 hours. For stable isotope labeled samples of WT N_175-245_ and R203M N_175-245_, BL21(DE3) *E. coli* with the appropriate plasmid were grown in MJ9 minimal media with ^15^N ammonium chloride and ^13^C glucose as the sole nitrogen or carbon source as appropriate. Cells were grown to an OD of 0.6-0.8 and induced with 1 mM IPTG at 37° C for 6 hours. Following induction, cells were harvested by centrifugation and either used immediately or stored at -80° C.

WT and R203M N_175-245_were purified using the TALON His-tag purification protocol (Clontech Laboratories). Cell pellets were lysed in high salt buffer (50 mM tris, 500 mM NaCl, 5 mM imidazole, 1 mM NaN_3_, pH 7.5) by sonication and centrifuged at 27200 RCF to remove all cell debris. The supernatant was mixed with resin for 1 hour and washed with high salt buffer (20X resin volume) and low salt buffer (50 mM tris, 150 mM NaCl, 5 mM imidazole, 1 mM NaN_3_, pH 7.5 (4x resin volume)). The SUMO solubility tag was cleaved with 200 nM SENP1 protease (∼1:250 protease:protein) for 1 hour at 4°C on the resin. GFP-N_175-245_was eluted off the resin with high salt buffer supplemented with 300 mM imidazole. The GFP-N_175-245_protein was exchanged into TEV cleavage buffer (50 mM Tris, 300 mM NaCl, 5 mM imidazole, pH 7.5) using a PD-10 desalting column (Cytiva). The GFP tag was cleaved by incubating overnight with 3 µM TEV protease (∼1:20 protease:protein) at 4° C, and then removed by reverse affinity purification with Talon His-Tag resin. Proteins were concentrated and exchanged into NMR buffer (50 mM sodium phosphate, 150 mM NaCl, pH 6.5).

Full-length SARS-Cov-2-Nucleocapsid (FLN) was prepared as previously described [3]. The R203M mutant plasmid of FLN (R203M-FLN) was generated by site directed mutagenesis. The plasmid was transformed into *E. coli* Rosetta (DE3) cells, cultured in 2xYT media to an OD of 0.6 and induced with 1 mM IPTG at 18° C overnight. Cells were harvested by centrifugation then resuspended in lysis buffer (50 mM sodium phosphate, 1 M NaCl, 5 mM imidazole, 1 mM NaN_3_, 1 mg/ml lysozyme, pH 8.0) and incubated for 1 hour at 4° C. Cells were sonicated 3x 2 minutes and centrifuged at 27200 RCF for 45 minutes. The clarified lysate was mixed with Talon His-Tag resin, incubated for 1 hour at 4° C, then washed with 20 column volumes of high salt FLN buffer (50 mM sodium phosphate, 3 M NaCl, 10 mM imidazole, 1 mM NaN_3_, pH 8.0). The protein was eluted with FLN elution buffer (50 mM sodium phosphate, 300 mM NaCl, 350 mM imidazole, 1 mM NaN_3_, pH 8.0), then concentrated and further purified using a Superdex 200 column in (50 mM sodium phosphate, 150 mM NaCl, pH 7.5) buffer. The protein was concentrated and either used immediately or stored flash frozen at -80° C.

pSer188 N_175-245_ plasmids were generated as previously described [31] and transformed simultaneously with pKW2-EFsep into BL21(DE3) ΔSerB *E. coli* cells. For natural abundance pSer188 N_175-245_, cells were grown in 2xYT to an OD of 0.6 to 0.8 and induced with 1 mM IPTG for 48 hours at 18 °C, then harvested by centrifugation at 2560 RCF. ^15^N-pSer188 N_175-245_, was expressed and purified as previously described [32]. Briefly, cells were grown to an OD 0.6 to 0.8 in MJ9 minimal media supplemented with ^15^N Celtone (0.2%), then induced with 1 mM IPTG for 48 hr at 18° C. Cells were harvested by centrifugation at 2560 RCF for 45 minutes and stored at - 80° C. Protein was purified using the same method as WT-N_175-245_, except with buffers supplemented with phosphatase inhibitors (10 mM NaF, 2.5 mM sodium pyrophosphate and 1 mM orthovanadate).

pSer188-FLN plasmid was generated by cloning the FLN sequence into the pRBC vector, then mutating S188 to an amber stop codon (TAG) for expression with genetic code expansion. The pSer188-FLN plasmid was transformed simultaneously with pKW2-EFsep into BL21(DE3) ΔSerB E coli cells. Cells were grown in rich-Auto-inducing Media [33] at 37° C until OD 600 reached 1.3, then at 18° C for 48 hours. Cells were harvested and the protein was purified using the same methods as FLN, except with buffers supplemented with phosphatase inhibitors (10 mM NaF, 2.5 mM sodium pyrophosphate and 1 mM orthovanadate). For hyperphosphorylated protein, 75 μM of target protein (pSer188 linker or FLN pSer188) was incubated with 80 nM of GSK3β in a buffer containing 20 mM Tris pH 7.4, 150 mM NaCl 10 mM MgCl_2_, and 1 mM ATP, at 37° C for 20 hours.

### NMR spectroscopy

NMR experiments were performed using an 800 MHz Bruker Avance III HD NMR spectrometer equipped with a triple resonance (TCI) cryogenic probe. Backbone resonance assignments were made using a suite of BEST triple resonance experiments, including HNCO, HNCACB, HNCACO, and HNCOCACB [34]. All NMR data were processed (apodized, zero filled, Fourier Transformed, and phased) using nmrPipe [35] and analyzed in nmrviewJ [36]. Three dimensional experiments were collected using non-uniform sampling (NUS) and NUS artifacts were suppressed using SMILE [37]. Resonance assignments were deposited in the BMRB under ascension number 51904. ^15^N nuclear spin relaxation parameters were measured using Bruker temperature compensated pulse sequences, with 8 unique delay times. The 60 ms (R_1_) and 34 ms (R_2_) delays were collected in triplicate to aid with error estimation. Peak intensities were fit in nmrviewJ to an exponential decay model, with Monte Carlo based error estimation. For the {^1^H}-^15^N NOE the D1 delay was increased to 8 seconds to ensure complete relaxation between scans. NMR titrations with RNA were performed using 2D ^15^N-BEST-TROSY experiments. Peak intensities were measured in nmrviewJ and normalized to the corresponding peak in the spectrum without RNA.

### Electrophoretic mobility shift assay (EMSA)

The first 1000 nucleotides from the viral genome (g(1-1000) RNA) at a final concentration of 0.5 μM was incubated with increasing concentrations of protein (range of 0-75 μM, in 20 mM Tris, 150 mM NaCl, 1 mM DTT, pH 7.5) at room temperature for 20 min. The total reaction volume was 10 µl. After incubation, 2 µL of 6X loading dye was added to the reaction before loading on a 1% agarose gel. The gel was run at 150 volts for 1 hour. RNA bands were stained with Midori Green Nucleic Acid staining solution (Bulldog Bio. Inc. Portsmouth, NH) and visualized using a Bio-Rad Gel Doc Image system.

### Sedimentation Velocity Analytical Ultracentrifugation

Sedimentation velocity analytical ultracentrifugation (SV-AUC) was performed using a Beckman Coulter Optima XL-A analytical ultracentrifuge equipped with absorbance optics. All SV-AUC experiments used linker protein tagged with GFP because the linker by itself does not have a strong UV absorbance. All samples were run in standard 2-channel sectored cells using an An60-Ti rotor. The concentration of each protein was varied from 50 to 200 μM. Samples were spun at 42,000 rpm and with 300 scans per sample. Data were fit to the continuous c(s) model using SEDFIT [38]. Buffer density and viscosity were calculated using SEDNTERP [39].

### LLPS and Microscopy

Fluorescence microscopy images were taken on a Keyence BZ-X700/BZ-X710 microscope with a 40X objective lens and a 384-well plate (Cellvis P384-1.5H-N); images were processed using BZ-x viewer and BZ-x analyzer software. For this experiment, cy3-labeled RNA was diluted into nuclease free water to reach a final concentration of 50 nM when added to the protein sample. Stocks of unlabeled protein were prepared by diluting into 20 mM Tris, 150 mM NaCl, 1 mM DTT, pH 7.5 droplet buffer. For FLN constructs, samples were prepared at a total concentration of 4 μM and 10 μM. For linker constructs, samples were prepared at a total concentration of 20 μM. Protein samples were prepared by combining 27 μL of protein stock of the appropriate concentration with 3 μL of cy3-labeled g(1-1000) RNA for a total sample volume of 30 μL. The samples were incubated at 37° C for 1.5 or 16 hours and subsequent imaging was taken. Foci were counted using ImageJ software. Average foci diameter was calculated in ImageJ from a random sampling of 10 foci from each image.

## Acknowledgments

We acknowledge helpful discussions from Richard Cooley, David Hendrix, and Brittany Lasher. We additionally acknowledge the support of the Oregon State University NMR Facility.

## Author contributions

Conceptualization: P.N.R., H.S., Z.Y., E.J.B. Methodology:P.N.R., H.S., Z.Y., S.S., K.H., E.J.B. Formal analysis: P.N.R., H.S. Investigation: P.N.R., H.S., Z.Y., S.S., K.H. Resources: H.S., Z.Y., S.S. Writing - Original Draft: P.N.R., H.S., Z.Y., S.S., K.H., E.J.B. Writing - Review & Editing: P.N.R., H.S., Z.Y., S.S., K.H., E.J.B. Visualization: P.N.R., H.S., E.J.B. Supervision: P.N.R., E.J.B. Project administration: P.N.R., E.J.B. Funding acquisition: E.J.B.

## Data and materials availability

All data needed to evaluate the conclusions in the paper are present in the paper and/or the Supplementary Materials.

## Supplementary Materials for

**Fig. S1.**
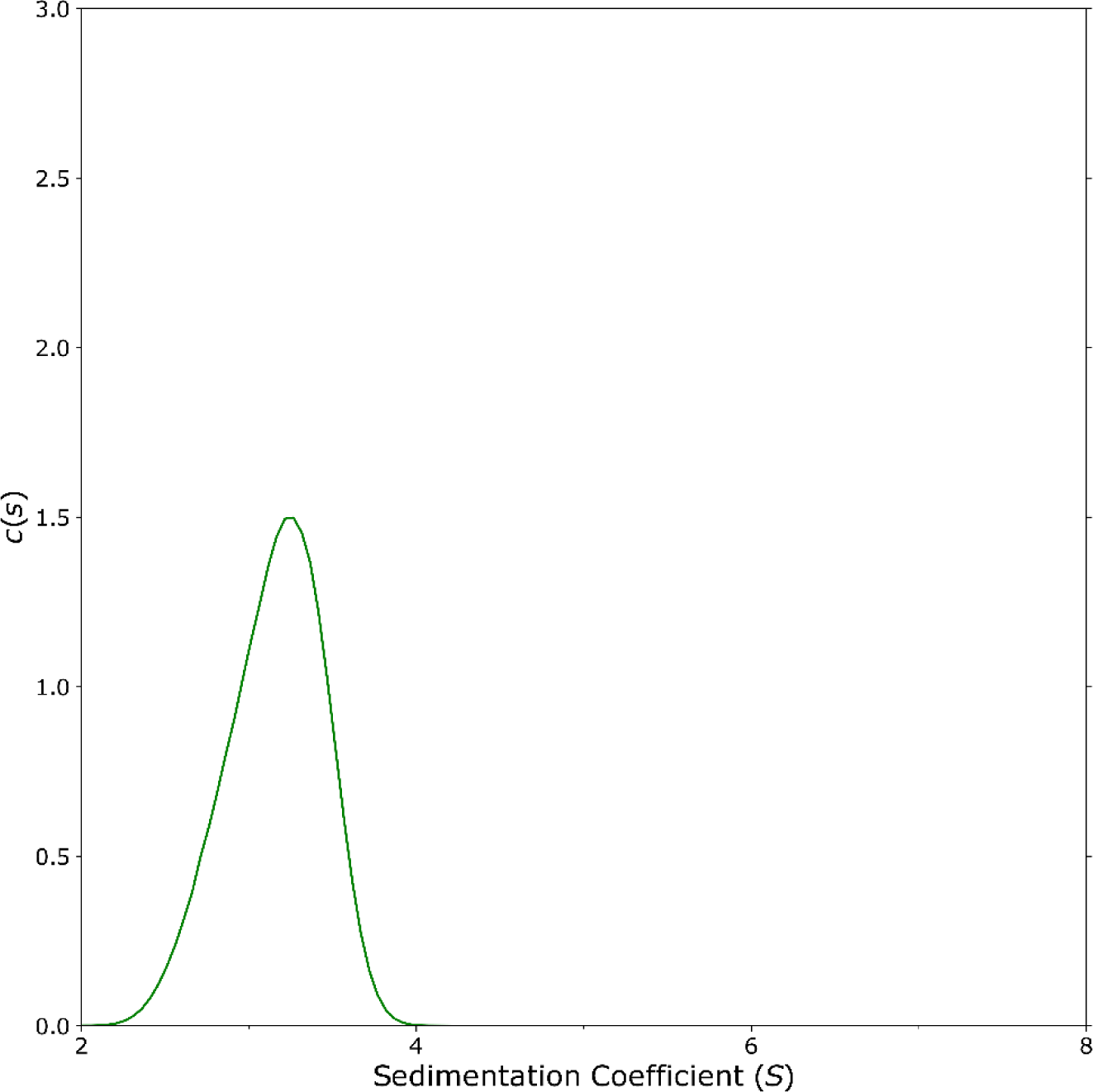
SV-AUC of GFP without N_145-245_. SV-AUC was performed on a sample of 200 μM GFP without N_145-245_. The observed sedimentation coefficient was ∼3.2 S.

**Fig. S2.**
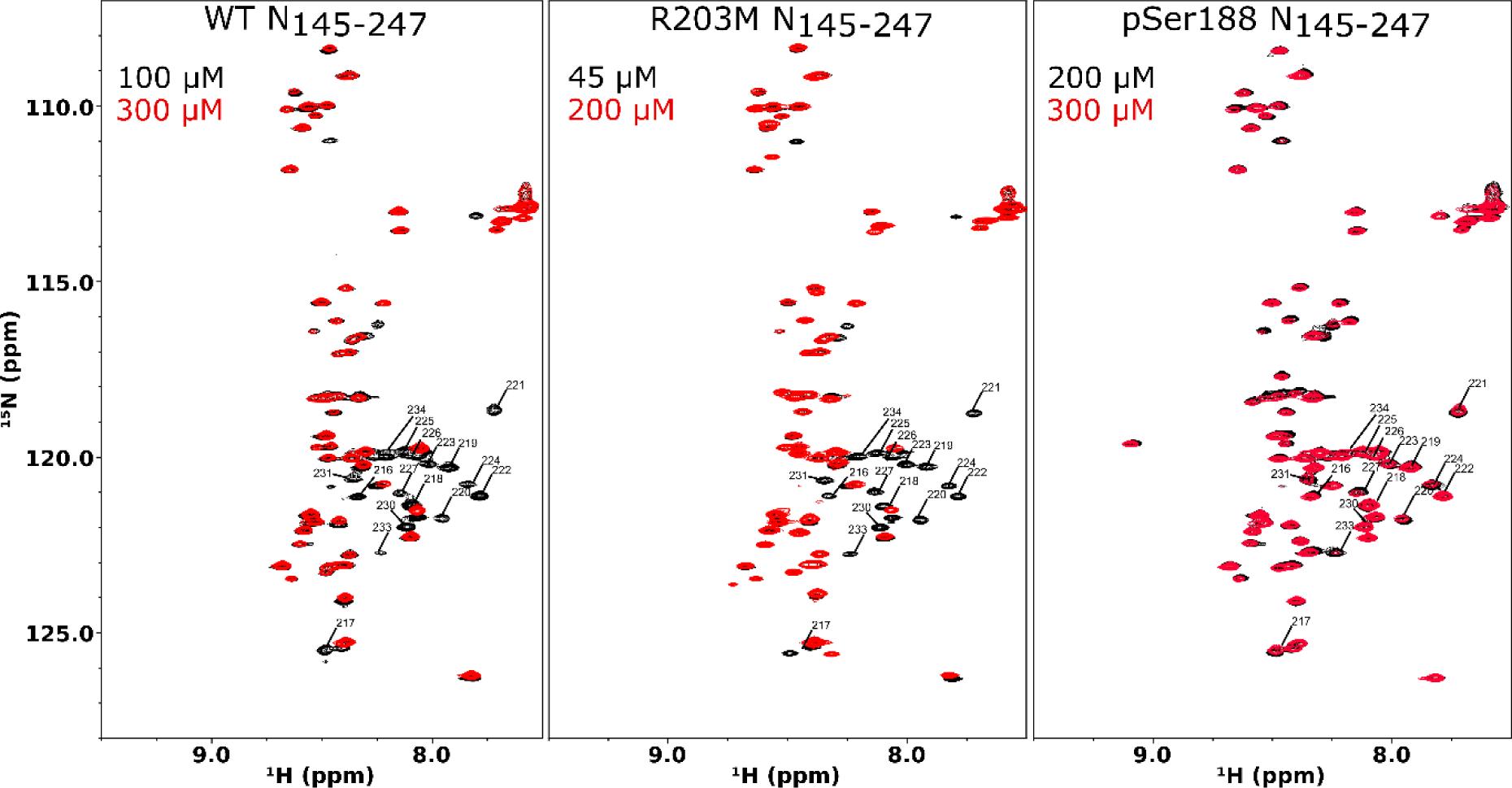
Comparison of N_145-245 15_N HSQC spectra of WT, R203M, and pSer188 variants at different concentrations. Higher concentration is shown in red, lower concentration is shown in black. Residues corresponding to the helical region are labeled.

**Fig. S3.**
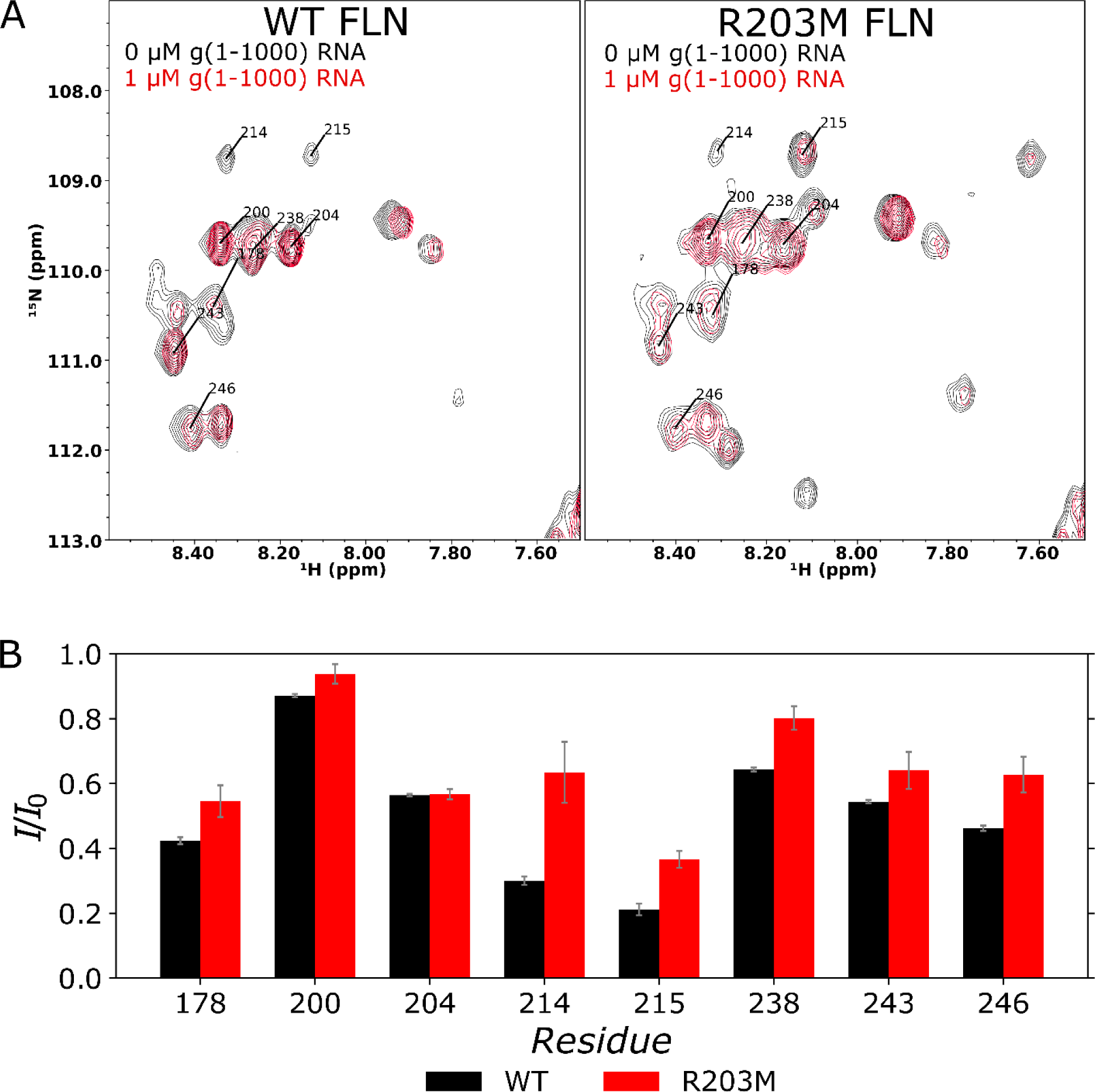
Comparison of WT and R203M FLN interaction with RNA. **A.** 15N HSQC spectra with 1 μM g(1-1000) RNA (red) and without RNA (black). Glycine residues assigned based on the N_145-245_ assignments are shown with labels. **B.** Comparison of peak intensities with and without RNA for the assigned glycine residues. The data are shown as the ratio of the intensity with RNA (I) over the intensity without RNA (I_0_). WT FLN is shown in black, R203M is shown in red. Error bars are one standard deviation, determined using the standard deviation of the noise in the spectrum.

**Fig. S4.**
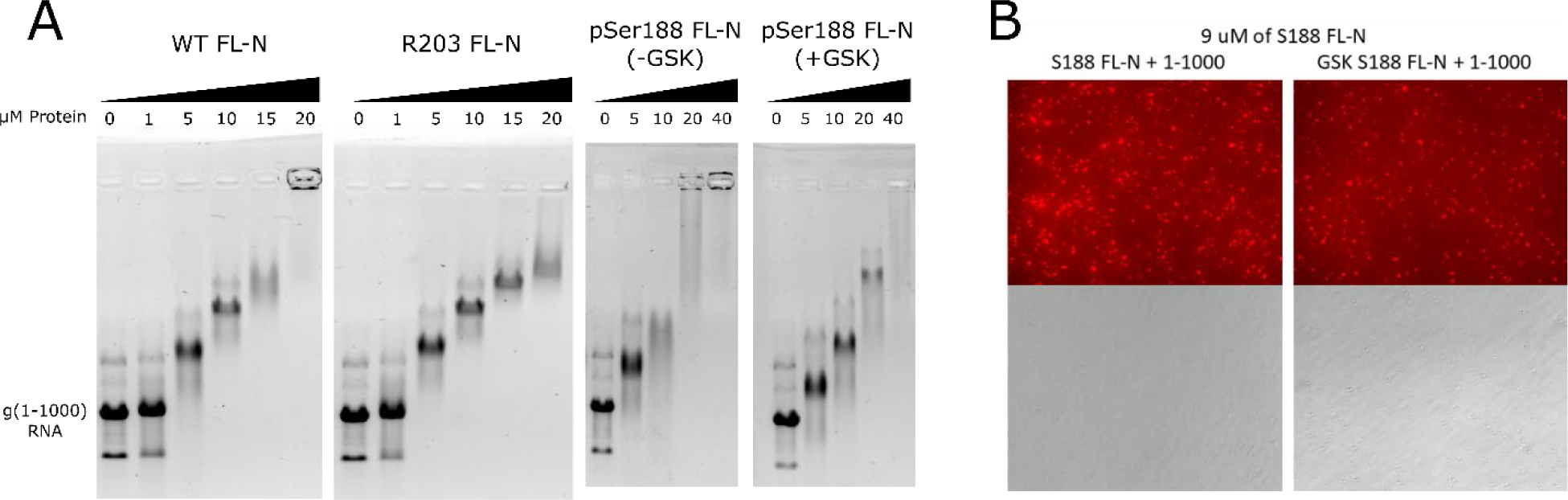
Interaction between RNA and FLN variant proteins. **A.** EMSA analysis of FLN binding to g(1-1000) RNA. For the phosphorylated FLN, EMSA was done with (+GSK) or without (-GSK) GSK-3. **B.** LLPS for pSer188 FLN at 9 μM protein.

